# A method to correct for local alterations in DNA copy number that bias functional genomics assays applied to antibiotic-treated bacteria

**DOI:** 10.1101/2023.07.10.548391

**Authors:** Geraldine J. Sullivan, Lars Barquist, Amy K. Cain

## Abstract

Functional genomics techniques, such as transposon insertion sequencing and RNA sequencing, are key to studying relative differences in mutant fitness or gene expression under selective conditions. However, certain stress conditions, mutations, or antibiotics can directly interfere with DNA synthesis, resulting in systematic changes in local DNA copy number along the chromosome. This can lead to artefacts in sequencing-based functional genomics data when comparing antibiotic treatment to an unstressed control, with relative differences in gene-wise read counts being the result of alterations in chromosomal replication dynamics rather than selection or direct gene regulation. We term this artefact ‘chromosomal location bias’ and implement a principled statistical approach to correct for it by calculating local normalization factors along the chromosome. These normalization factors are then directly incorporated in statistical analyses using standard RNA-sequencing analysis methods without modifying the read counts themselves, preserving important information about the mean-variance relationship in the data. We illustrate the utility of this approach by generating and analysing a ciprofloxacin-treated transposon insertion sequencing dataset in *Escherichia coli* as a case study. We show that ciprofloxacin treatment generates chromosomal location bias in the resulting data, and we further demonstrate that failing to correct for this bias leads to false predictions of mutant drug sensitivity as measured by minimum inhibitory concentrations. We have developed an R package and user-friendly graphical Shiny application, ChromoCorrect, that detects and corrects for chromosomal bias in read count data, enabling the application of functional genomics technologies to the study of antibiotic stress.

**IMPORTANCE:** Altered gene dosage due to changes in DNA replication has been observed under a variety of stresses with a variety of experimental techniques. However, the implications of changes in gene dosage for sequencing-based functional genomics assays are rarely considered. We present a statistically principled approach to correcting for the effect of changes in gene dosage, enabling testing for differences in the fitness effects or regulation of individual genes in the presence of confounding differences in DNA copy number. We show that failing to correct for these effects can lead to incorrect predictions of resistance phenotype when applying functional genomics assays to investigate antibiotic stress, and we provide a user-friendly application to detect and correct for changes in DNA copy number.

## Introduction

The capacity to associate genes with their functions is a critical driver of scientific progress. This is reflected in the 2023 U.S. Biotechnology and Biomanufacturing policy, which has set the goal of sequencing of 1,000,000 diverse microbes and determining the function for 80% of newly discovered genes (1). Functional genomics technologies, such as transposon insertion sequencing (TIS) (2) and RNA-sequencing (RNA-seq) (3), have emerged as effective, high-throughput methods for investigating gene function and will be instrumental to assigning function to new genes *en masse*. Currently, most functional genomics methods rely on quantifying and comparing sequencing read counts, with results often expressed as relative (log) ratios of read counts between an experimental condition and control. For instance, the TIS technique involves generating a large pool of random transposon mutants which can be uniquely identified by selectively amplifying and sequencing the flanking chromosomal DNA using transposon-specific primers. Then TIS analysis pipelines such as the TraDIS toolkit (4) or MAGenTA (5) assess changes in mutant populations with or without selection, using transposon insertion read counts per gene as proxies for estimating mutant abundance. The typical output of a TIS analysis is a log_2_ fold change (log2FC) in mutant abundance for each genetic feature, generally interpreted as reflecting the relative fitness of the mutant. Similarly, differential expression analysis of RNA-seq uses read counts to generate a log2FC per gene, which represent changes in relative RNA expression levels.

Since their initial discovery nearly a century ago, a wide variety of antibiotics have been developed and traditional laboratory studies have elucidated their primary antimicrobial mechanisms and targets for many bacterial species (6, 7). For example, certain antibiotic classes, such as quinolones, are known to interfere with the process of DNA replication by inhibiting topoisomerases, leading to cell death (8). However, further molecular investigations of commonly used antibiotics are still underway, aiming to fight against the rising threat of antibiotic resistance and aid further antibiotic development. Detailed genomics-based studies have uncovered that antibiotics can exhibit signs of secondary targets and knock-on effects that could be exploited as helper drug targets (9–12). For example, ciprofloxacin can also affect filamentation or outer membrane composition (13, 14). These studies can also aid in tracking bacterial responses to antibiotic exposure as well as understanding how resistance is gained, predicted, and potentially prevented (15–17). Many studies have employed functional genomics techniques such as TIS and RNA-seq to comprehensively study antibiotic response and determine each gene’s contribution to cell survival during antibiotic exposure for a wide-range of bacterial species and antibiotic classes. For instance, L. A. Gallagher et al. (18) identified novel isomerases that aided in tobramycin tolerance using TIS in *Pseudomonas aeruginosa*, B. Jana et al. (19) identified the secondary resistome of *Klebsiella pneumoniae* for the last-line polymyxin colistin, C. J. Boinett et al. (20) utilized TIS and RNA-seq to identify multiple resistance responses to colistin treatment in *Acinetobacter baumannii* (20), and others have subjected bacteria to panels of antibiotics to compare cross-antibiotic responses (21–23).

During normal exponential growth in bacteria, the time between cell divisions is significantly shorter than the time needed to complete chromosomal replication. This leads to cells containing more copies of DNA around the origin of replication (*oriC*) compared to the terminus (*ter*) and is due to the firing of multiple simultaneous replication forks (24) (**Figure 1A**). In many sequencing-based assays, this difference in DNA copy numbers translates into higher read counts around the origin as compared to the terminus due to a higher availability of template nucleic acids. Under most conditions, the ratio of *oriC-ter* reads remains constant when performing differential analysis between a treatment and an untreated control and does not interfere with results. However, conditions that specifically alter the *oriC-ter* ratio, such as certain antibiotic treatments like ciprofloxacin, may introduce large distortions in read counts (**Figure 1B**), leading to fold changes that primarily reflect differences in the local chromosomal copy number (**Figure 1C**). The impact of altered local copy numbers on read counts and subsequent fold changes in differential analysis is what we have termed ‘chromosomal location bias’ (CLB). Previously, a comprehensive study by Slager and colleagues (25) showed that treatment with some or all of a range of antibiotics and antimicrobials (ciprofloxacin, trimethoprim, hydroxyurea, 6(p-Hydroxyphenylazo)-uracil (HPUra), N3-hydroxybutyl 6-[3’-ethyl-4’-methylanilino]-uracil (HBEMAU), mitomycin C) modified both DNA and RNA copy number, resulting in altered *oriC-ter* ratios across the genomes for multiple bacterial species including *Streptococcus pneumoniae*, *Bacillus cereus*, *Staphylococcus aureus* and *E. coli* (25). In a subsequent review, Slager and colleagues (26) suggested that not normalizing for this effect in RNA-seq analyses may lead to an overestimation of differential gene expression.

**Figure 1.**
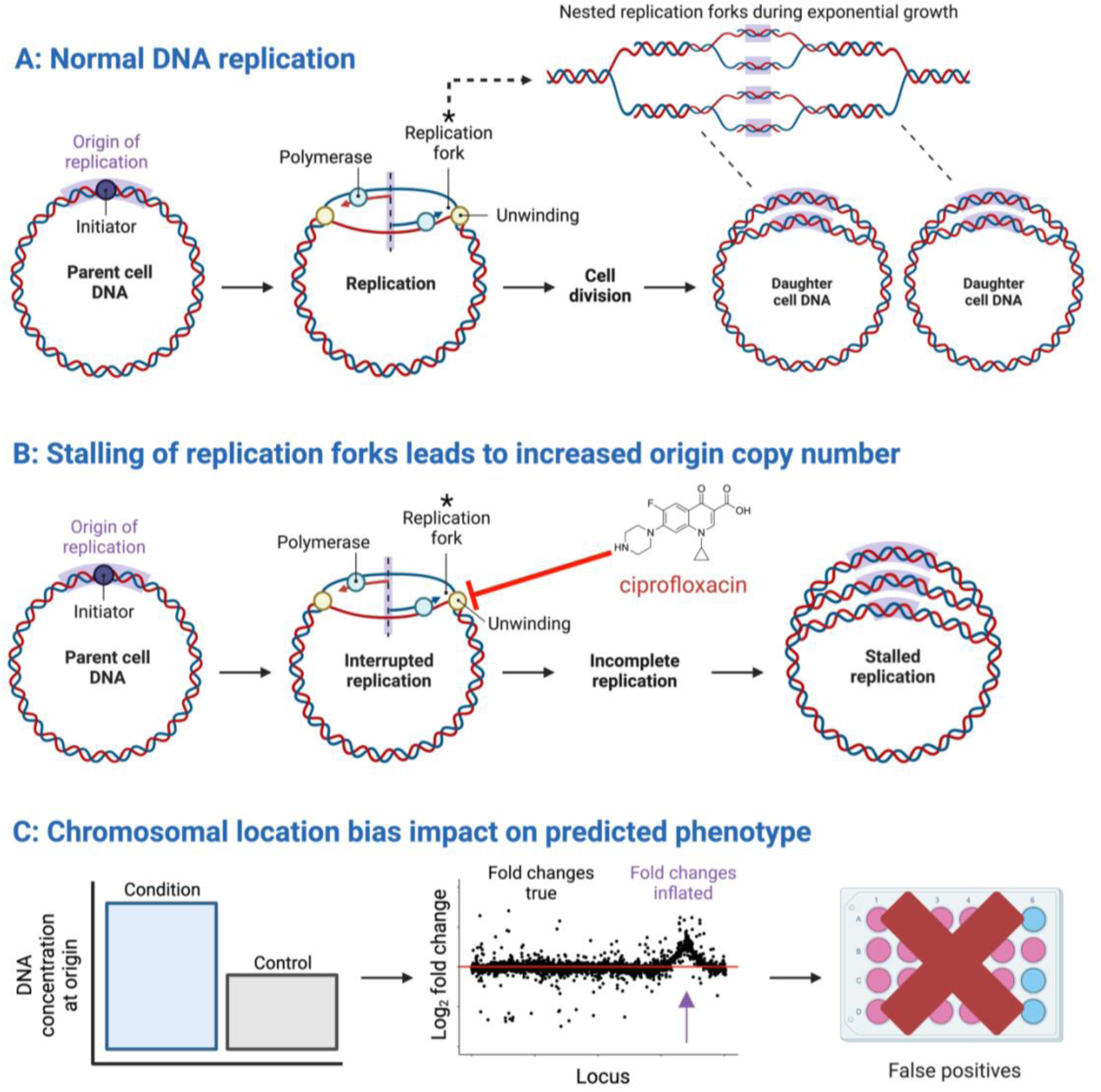
Chromosomal location bias as a result of higher chromosomal copy number near the origin after ciprofloxacin treatment and the downstream effects on read counts. A) Normal DNA replication producing two daughter cells. The origin of replication is colored purple. Multiple replication forks naturally lead to more origin (*oriC*) than terminus (*ter*) DNA. B) Ciprofloxacin prevents DNA unwinding, stalling the replication fork. This produces a highly inflated relative concentration of DNA proximal to the origin. C) The knock-on effects of increased DNA concentration around the origin on the read counts and fold changes, an observable peak at the origin in the relative log2FCs and potential false positive predictions of downstream drug sensitivity testing.

The consequences of CLB for interpretation depends on the functional genomics technology employed. In the case of RNA-seq, CLB is reflective of a genuine increase in RNA synthesized from the origin-proximal region, and the primary danger is that this could be interpreted as a specific regulatory response rather than a direct result of antibiotic activity. As differential expression is calculated with respect to the average count ratio between experiments, CLB may also mask regulated expression if the changes in expression are smaller than the changes induced by differences in DNA copy number. For TIS experiments, CLB introduces artefacts that can lead to false predictions of phenotype. Since TIS uses transposon-flanking reads as a proxy for mutant abundance, and local distortions in DNA copy number will lead to a local distortion in template DNA abundance, mutants containing transposon insertions in the vicinity of the origin will appear to be more frequent in the population than they really are in data affected by CLB. This can lead to false predictions of drug sensitivity or resistance if not corrected for.

Previous studies have attempted to correct CLB in antibiotic-treated functional genomics data using local regression methods such as Lowess (18, 23, 27). Typically, local regression is performed on the raw read counts, with normalized counts produced to replace the raw reads for the differential analysis. However, most differential analysis tools for sequencing data rely on count models that assume counts of similar magnitude have similar variance (28, 29). Providing modified or transformed counts violates these assumptions and will lead to incorrect assessment of statistical significance. We often observe distortions in the local read count density spanning several orders of magnitude following antibiotic treatment, which would lead to concomitantly large distortions in resulting p-values.

Here, we develop a statistically principled approach for correcting CLB that incorporates a local normalization factor as an offset in edgeR (28), rather than directly providing normalized counts. This preserves important features of the data needed for accurate calculation of p-values, namely the mean-variance trend, while also producing fold-changes that have the CLB effect removed. Based on this method we have developed an application for identifying and correcting for CLB named ‘ChromoCorrect’. We have made our diagnostic and normalization procedure availably as a graphical Shiny application that can be applied to any sequencing based functional genomics assay. We apply ChromoCorrect to a dataset we generated for this study that displays strong CLB: TIS output of an *E. coli* K12 library challenged with ciprofloxacin. We confirm that ciprofloxacin produces the predicted large local distortions in read counts around the origin of replication, confounding the TIS counts such that they no longer accurately reflect the fitness of individual mutants. We show using minimum inhibitory concentration (MIC) assays that these distortions lead to incorrect predictions of mutant ciprofloxacin sensitivity and resistance. We also demonstrated that our normalization approach, after processing with ChromoCorrect, corrects these, rendering accurate fold changes that align well with independently determined mutant phenotypes.

## Results

### Chromosomal location bias distorts a range of functional genomics datasets

To illustrate the prevalence of CLB, we have collected instances of CLB in functional genomics data resulting from a variety of treatments in multiple organisms (**Figure 2**). Slager and colleagues (25) showed that the uracil analogue HPUra induced competence in *S. pneumoniae* as an indirect effect of stalling replication forks, increasing the *oriC-ter* ratio and thereby increasing the expression of origin-proximal genes (25). They demonstrated a clear instance of CLB by plotting RNA-seq log2FCs from treatment with HPUra and untreated bacteria along the chromosome (**Figure 2A**). For the current study, we additionally collected various published functional genomics data sets representing a variety of antibiotic classes and bacterial species and then examined them for CLB.

**Figure 2.**
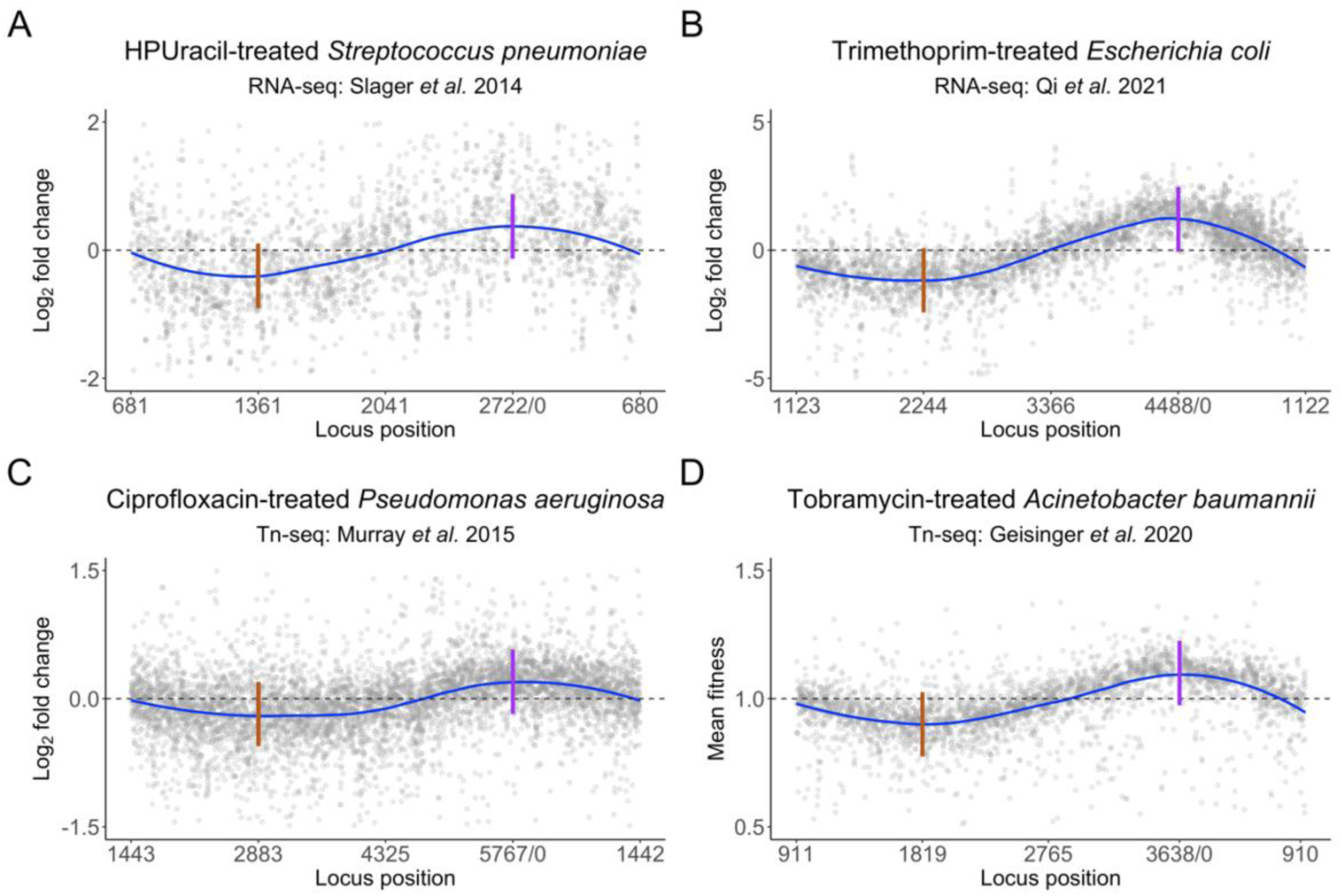
Read count log_2_ fold changes vs genome location plots displaying chromosomal location bias under varying conditions and organisms using different experimental techniques. Blue lines indicate trendlines over the individual grey points. Dashed grey lines indicate the expected trend line. Solid vertical lines indicate the terminus (brown) and origin of replication (purple). All experiments have decreased fold changes or fitness scores that dip at the terminus and increase at the origin. A) RNA-seq analysis of 6(p-Hydroxyphenylazo)-uracil-treated *Streptococcus pneumoniae*. B) RNA-seq analysis of trimethoprim-treated *Escherichia coli*. C) Tn-seq analysis of ciprofloxacin-treated *Pseudomonas aeruginosa*. D) Tn-seq analysis of tobramycin-treated *Acinetobacter baumannii*. Data sources: A: Slager *et al*. (25), B: Qi *et al*. (30), C: Murray *et al*. (21) D: Geisinger *et al*. (23).

Firstly, we extracted RNA-seq data from a study looking at *E. coli* treated with subinhibitory concentrations of trimethoprim (30), an antibiotic commonly used in the treatment of urinary tract infections (31). Trimethoprim targets the dihydrofolate reductase of the folate biosynthesis pathway and reduces availability of tetrahydrofolate, a precursor to the essential DNA components thymidine and thymine (32) eventually leading to so-called thymineless death (33). Thymine starvation stalls replication forks, initially leading to a transient increase in origin-proximal DNA before the replicating DNA is destabilized and degraded, leading to ultimate depletion of origin-proximal DNA (34). As would be expected for subinhibitory treatment, we observe inflated fold changes in the vicinity of the origin (**Figure 2B**).

The presence of CLB in TIS data could potentially be a more serious confounder of downstream interpretation than in RNA-seq as TIS amplifies mutant DNA directly as a proxy for mutant cell frequencies. Thus, any condition-specific change in local DNA concentration within individual cells will translate into distorted fold changes. We collected two sets of TIS data illustrating that this effect occurs across organisms and antibiotic classes. We examined Tn-Seq data from *Pseudomonas aeruginosa* treated with ciprofloxacin (21), which we predict to harbor CLB (**Figure 1B**) and when plotted, it indeed showed clear evidence of a local increase in DNA copy number around the origin (**Figure 2C**). Further, CLB had previously been observed in *P. aeruginosa* treated with the ribosome-targeting antibiotic tobramycin (18). Therefore, we collected data from a recent study applying Tn-seq to study the effects of tobramycin treatment on *A. baumannii,* and it showed the characteristic CLB wave pattern (**Figure 2D**) (23). Taken together, we surmised that CLB may be relatively wide-spread in antibiotic-treated functional genomics datasets and that a robust method to detect and remove CLB would be beneficial for researchers to produce accurate predictions for downstream analysis.

### A principled normalization procedure to correct for chromosomal location bias

To correct for CLB, we have developed a normalization procedure that produces offsets that can be directly incorporated into differential testing using edgeR (28) or DEseq2 (29) without modifying the input count data. These offsets can be thought of as a gene-specific normalization factor, which in this case includes a correction for the local read density across the chromosome. A similar approach has been taken by the transcript quantification package tximport to correct for differences in isoform abundance during gene-level differential expression testing in eukaryotes (35).

Our normalization procedure comprises three major steps as outlined in **Figure 3**. First, we calculate the local median read depth along the chromosome using a sliding window over the local gene neighborhood. The number of flanking genes included in the sliding window starts at 500 and is dynamically determined from the data set by fitting a linear model and testing the slope and y-intercept of the fitted line. Each iteration reduces the window size by 100 loci if the window is not small enough to accurately fit the trendline until a minimum window size of 200 is reached. Although the median is more robust at handling outliers compared to the mean, we have still found that medians calculated from windows smaller than 200 loci can be unduly influenced by a small number of genes with particularly high or low read counts. Second, the sliding window analysis is used to calculate a gene read count normalized for local read density by dividing the actual read count for each gene by the ratio of the local to global average read counts. This normalized count is then used to derive a gene-wise normalization factor, additionally incorporating differences in effective library size between replicates using the trimmed mean of M-values (TMM) method (36) (see Methods). Finally, these offsets are provided to the edgeR (28) glmFit function alongside the raw counts for differential analysis, where they are directly incorporated into the statistical model for testing purposes. This procedure maintains the information contained in the raw counts, necessary for accurate statistical analysis, while correcting the resulting estimated log2FCs for local distortions in DNA copy number.

**Figure 3.**
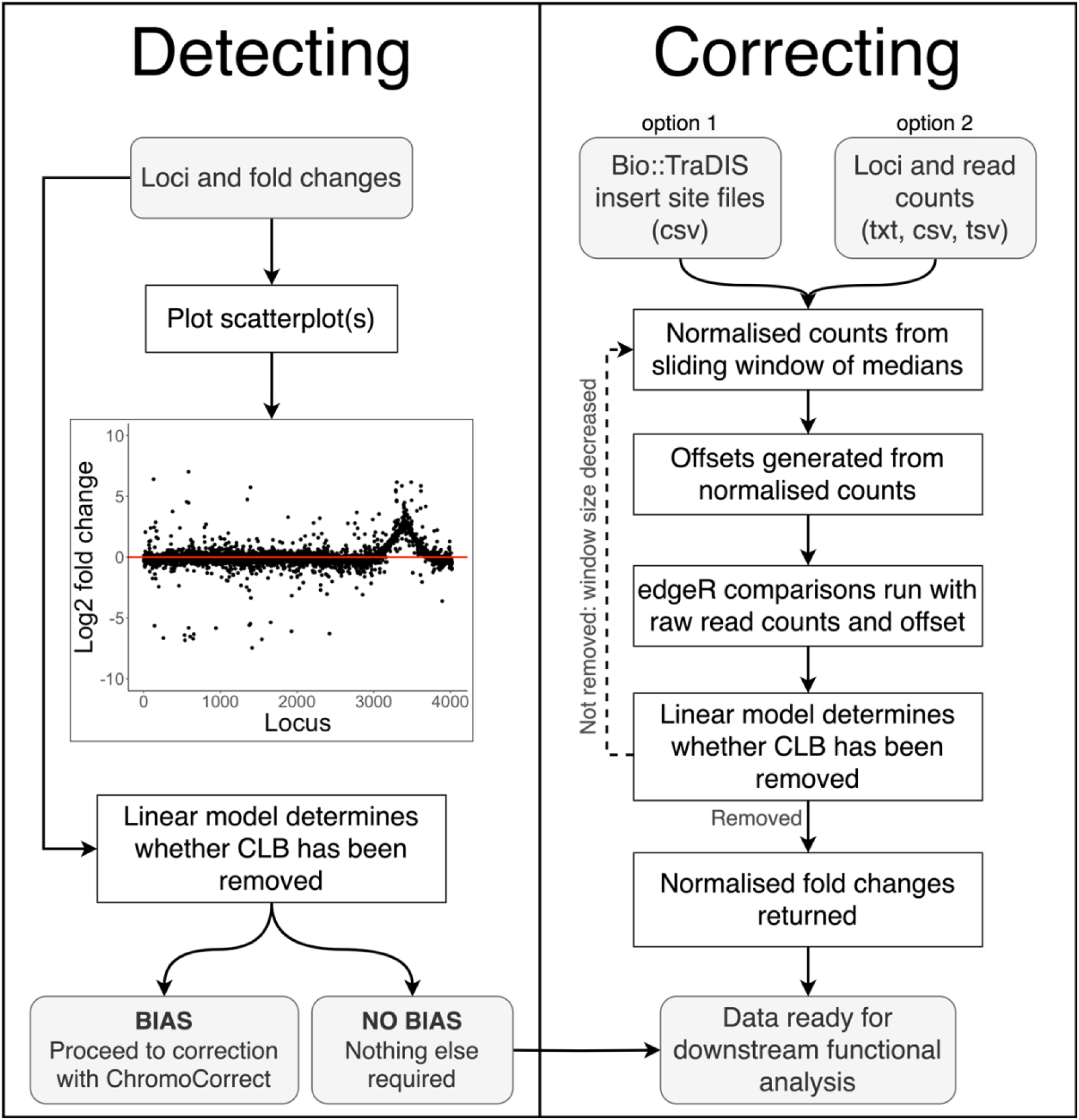
Schematic of pipeline for detecting and correcting chromosomal location bias using ChromoCorrect. Grey boxes indicate user input or output, and white indicates automated steps. Detecting requires the log_2_ fold changes of each locus to plot a scatterplot of fold change by chromosomal position, which graphs trends in the fold changes for the user to visualise the pipeline’s assessment. The app and the R console display a message recommending normalization if chromosomal location bias is detected by a fitted linear model. Correcting requires read counts per locus in a txt, csv or tsv file format, which are then normalized using a sliding window of medians with a default size of 500. An offset matrix is generated from the normalized counts to input along with the raw read counts into edgeR. A linear model is fitted again during correction to determine whether the default sliding window is small enough to capture the trend and repeats the normalization procedure with a smaller window otherwise. The corrected analysis is returned after the normalization is complete.

### The ChromoCorrect app

The normalization techniques described here have been organized into an R Shiny app for easy use by researchers wanting to diagnose and normalize data affected by CLB. Instructions for installing and running the app can be found on Github (https://github.com/BarquistLab/ChromoCorrect/) or accessed online through ShinyApps (https://thecainlab.shinyapps.io/ChromoCorrect/). The app is split into two main tabs: *detecting* and *correcting* (**Figure 3**). *Detecting* requires the upload of analyzed output files containing log2FC values to visualize any CLB, whilst the *Correcting* tab requires upload of the read counts for the conditions affected by CLB and the associated no-stress control.

The first step within the app is to assess whether data sets that have undergone differential analysis are affected by CLB. This can be done using the *Detecting* tab. The user inputs one or more files containing locus tags and fold change information, and a locus by fold change scatterplot for each condition is generated. The user can cycle through the uploaded data sets in the drop-down menu in the sidebar to determine if any of the experimental conditions are affected by CLB. It is deemed present if the general trend of the fold changes is not flat and distributed around zero, as demonstrated in **Figure 3A**. The ‘decision’ box will contain red text suggesting correction if CLB is detected, or green text if not detected.

If the analysis produced CLB, the user can fix the issue with the second tab: *Correcting*. This tab requires the upload of a file with a locus_tag column and read counts, or the upload of read files produced by the tradis_gene_insert_sites script from Bio-TraDIS (4). The tab requires at least four insert site files or columns of read counts: two biological replicates of the experimental condition and two biological replicates of the associated control condition. Once the upload is complete, the user must define which condition specifies the control. The app will compute an edgeR comparison before and after normalization and produce two scatterplots of locus versus log fold change associated with the two analyses. The default window size of 500 is suitable for smooth trend lines. The window will automatically reduce in size if the data has not been normalized effectively due to any sharper trend lines present. The user can then export the corrected output, free from CLB.

### Transposon insertion sequencing case study: ciprofloxacin

To illustrate the functionality of ChromoCorrect, we generated a data set applying the transposon-directed insertion-site sequencing (TraDIS) TIS technique to a dense library of *E. coli* transposon mutants exposed to ciprofloxacin compared to an untreated control. This dataset was generated using *E. coli* K12 BW25113, the parental strain of the Keio collection (37). A TraDIS library of 350,000 unique Tn*5* mutants (38) was challenged with a subinhibitory concentration of ciprofloxacin (1/2 MIC) with growth overnight. After analysis with the TraDIS toolkit (4), we confirmed ciprofloxacin as an inducer of exaggerated CLB as it displayed a distinctive peak of increased reads around the origin of replication (**Figure 4A**). This peak reflects the expected increase in DNA copy number at the origin compared to the rest of the genome that ciprofloxacin induces as it targets topoisomerases and stalls the replication fork and DNA synthesis (25, 39). Around the peak, the observed inflation in transposon insertions occurred largely between loci 3023 and 3622, 300 loci on either side of the origin of replication (*oriC* located between locus 3322, *mnmG* and locus 3323, *mioC*) . This entire 600 locus region had an average log2FC of 1.5 (a fold change increase of 2.8 compared to the untreated control), whereas the rest of the genome had a more typical average log2FC of 0.1. Strikingly, this means that many of these 600 loci would meet the standard ‘2-fold’ cut-off often used to prioritize genes for further investigation.

**Figure 4.**
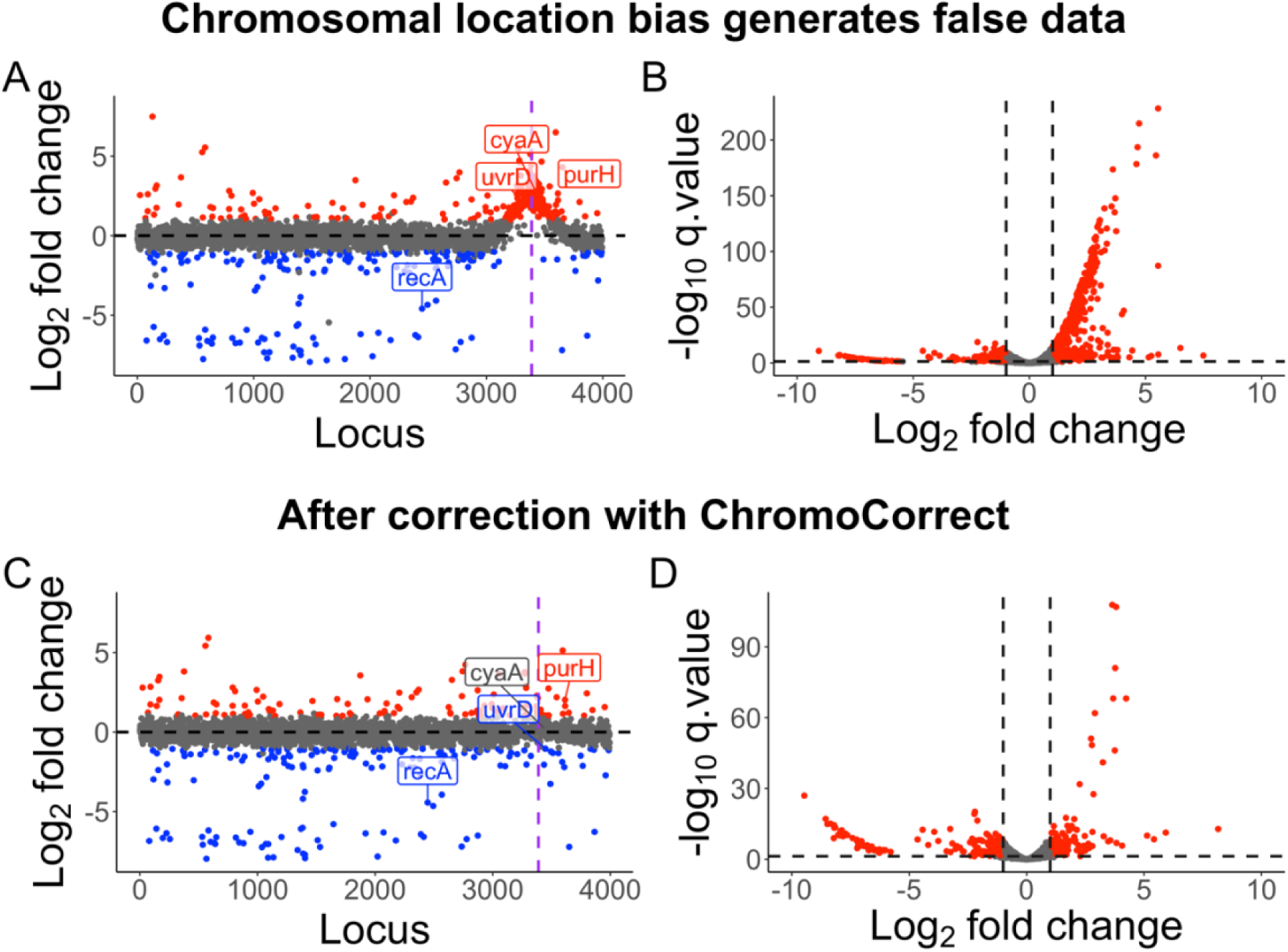
Visualizing and diagnosing chromosomal location bias in a ciprofloxacin-treated transposon insertion sequencing data set. Each point represents a locus of the genome. The x-axis is the chromosomal location, and the y-axis is the log2FC from ciprofloxacin treated versus untreated comparisons. A) Volcano plot before correction, showing a large skew of significant genes (red) to the right, representing an increased prevalence of these mutants compared to the no antibiotic control. B) Locus by fold change scatterplot pre-normalization with loci plotted in chromosomal order. The dashed black line shows the expected trend of the data if not affected by CLB. The purple dotted line is the origin of replication, where a large peak of elevated read counts is seen. C) Locus by fold change scatterplot post normalization with no peak. D) Volcano plot post-normalization with no skew. Normalization performed with a sliding median window size of 200. Some mutant examples from our phenotypic validation are labelled in the scatterplots. Blue genes represent significant genes with log2FCs ≤ -1, and red genes indicate significant genes with log2FCs ≥ 1.

Diving deeper into the data, the ciprofloxacin data after analysis with the TraDIS pipeline has 754 significant genes (q-value < 0.05), with 468 having an absolute log2FC value over 1. Of these 468 genes, 391 (84%) of these were located between loci positions 3023 and 3623, despite this region representing only 15% of the genome. Another visualization confirming this data bias was the volcano plot showing log2FC vs -log_10_ q-value (**Figure 4B**). The fold change was skewed to the right, indicating a bias towards mutants with a higher frequency of insertions in many genes. After normalizing the data using ChromoCorrect with an automatically determined sliding window median of 200 genes, we identified 272 significant genes (a 64% reduction) with only 163 having an absolute log2FC value over 1 (a 65% reduction). The normalized volcano plot and locus by fold change scatterplot shows the CLB has been removed (**Figures 3C** and **3D**).

### Chromosomal location bias leads to incorrect predictions of ciprofloxacin sensitivity

Using TIS or other functional genomics techniques to identify entire gene suites involved in antibiotic stress tolerance is well-established (18, 21, 22). However, the possible presence of CLB in these datasets, if left uncorrected, may result in false positives that are carried on into laboratory analysis. To demonstrate the implications of CLB for predictions of antibiotic resistance (or sensitivity), as well as to validate the use of ChromoCorrect to generate more biologically accurate predictions, we tested the phenotypes of various *E. coli* BW25113 single gene mutants the Keio collection (37) that represent different types of potential errors that may arise because of CLB. In total, 11 mutants were tested for their ciprofloxacin resistance and sensitivity profile (between 5-20 ng/ml) via a minimum inhibitory concentration (MIC) assay and compared to the wild type (WT) BW25113 resistance level (10 ng/ml) (**Table 1**). For this, we examined genes that exhibited dissimilar outputs before and after correction, particularly focusing on those that were identified as significant prior to correction but not afterwards.

**Table 1.** Single gene *E. coli* BW25113 Keio knockouts validated in this study.

| Predicted Mutant Phenotype | Gene | Function | TraDIS log2FC |  | Experimental Mutant Phenotype MIC<sub>50</sub> (ng/mL) |
| --- | --- | --- | --- | --- | --- |
|  |  |  | Before ChromoCorrect | After ChromoCorrect |  |
| - | WT | - | - | - | 10 |
| Cip sensitive before and after | *acrB* | multidrug efflux system protein | -0.89 | -0.83 | ≤5 |
|  | *recA* | DNA recombination and repair protein | -1.99 | -1.92 | ≤5 |
| Cip resistant before and WT phenotype after | *cyaA* | adenylate cyclase | 2.48 | 0.15 | 10 |
|  | *ilvN* | acetolactate synthase 1 small subunit | 2.13 | -0.06 | 10 |
|  | *mtlD* | mannitol-1-phosphate dehydrogenase | 1.77 | 0.13 | 10 |
|  | *fdhE* | formate dehydrogenase formation protein | 1.66 | 0.13 | 10 |
|  | *pfkA* | 6-phosphofructokinase I | 1.32 | -0.03 | 10 |
| Cip resistant before and after | *purH* | bifunctional transformylase/cyclohydrolase | 2.50 | 1.64 | 20 |
|  | *dgkA* | diacylglycerol kinase | 1.62 | 1.07 | 20 |
|  | *rpe* | D-ribulose-5-phosphate 3-epimerase | 1.45 | 1.07 | 20 |
| Cip resistant before and sensitive after | *uvrD* | DNA-dependent ATPase I and helicase II | 1.32 | -1.06 | ≤5 |

First, to sanity-check our normalization process and MIC assay, we checked the phenotype of genes known to play a role in ciprofloxacin tolerance. We validated the known antibiotic resistance determinants *acrB*, involved in efflux (40, 41), and *recA*, involved in DNA recombination and repair (42, 43). As expected, both genes remained significant after correction with ChromoCorrect with a predicted sensitivity phenotype, i.e., the genes are needed for ciprofloxacin tolerance. Further, our phenotypic mutant validation confirmed their importance in ciprofloxacin resistance, as both gene mutants showed a decreased MIC_50_ (≤5 ng/mL) compared to WT (10 ng/mL).

Next, we focused on genes where CLB led to a predicted sensitivity phenotype (with apparent increased mutant abundance) before but not after applying ChromoCorrect. Of 3920 analyzed genes, 543 (14%) were not significant after correction that were classified as significant before. Of this group, 335 met the typical phenotypic follow up threshold of an absolute log2FC ≥ 1 before correction, all of which had increased insertions and were located adjacent to the origin. Five genes previously unlinked to antibiotic resistance were chosen for the phenotypic validation based on availability of Keio mutants.

As predicted by the output of ChromoCorrect, all 5 mutants displayed no change in MIC compared to WT, confirming that without correction, these were falsely designated as ciprofloxacin sensitivity genes. We then tested three gene representatives that remained significantly more abundant after correction, and all showed a two-fold increase (20 ng/ml) in ciprofloxacin MIC, confirming a true sensitivity phenotype.

Lastly, we examined genes with the most extreme mispredictions of phenotype, those that shifted from predicted mutant resistance to predicted sensitivity after correction. There were eight genes whose insertion count went from positive to negative values, but only one (*uvrD)* met the phenotypic validation threshold of log2FC ≥ 1 before and after correction. There were no instances of significant predicted mutant sensitivity changed to resistance phenotypes due to the right-hand skew of the data (as shown in **Figure 4B**). The misprediction of *ΔuvrD* phenotype prior to correction was confirmed by the MIC validation, where the gene deletion was shown to have increased sensitivity. Neglecting to address this bias would have falsely classified *uvrD* as mediating ciprofloxacin sensitivity, not resistance. Overall, all mutants tested by MIC assay reflected the predicted phenotype after correction of CLB, validating its use in normalizing this data. This emphasizes the critical role of ChromoCorrect’s normalization in ensuring accurate and reliable gene fitness assessments.

## Discussion

This study addresses the fundamental issue of CLB, which we show impacts a wide variety of functional genomics analyses, resulting in false positives and negatives or incorrect interpretation. CLB arises from nucleic acid copy number fluctuations along the chromosome, typically around the origin and/or terminus, which can be exacerbated by replication disrupting events like replication-targeted antibiotic treatment. To solve this problem, we introduce ChromoCorrect, a normalization tool that effectively corrects for CLB producing accurate log2FCs and significance values for each locus. Using a ciprofloxacin-treated TraDIS dataset in *E. coli*, we demonstrated that ChromoCorrect corrects for CLB between an experimental condition and a control and leads to a more accurate prediction of antibiotic resistance phenotype.

It should be noted that for TIS experiments where mutant abundance is determined, correcting for CLB is essential to determine the true mutant fitness brought about by the comparison condition. However, for RNA-seq, the uncorrected data does reflect a genuine difference in RNA copy numbers which likely has an impact on the cellular response. While the correction method presented here will work on RNA-seq data and any experiments generating read counts, the decision on whether to perform the correction will depend on what questions the experiment is attempting to answer.

By employing correction with ChromoCorrect, we can collectively ensure that future differential functional genomics experiments accurately report the true nature of genes in conditions producing CLB, minimizing the reporting of false positives and advancing our understanding of gene function.

## Materials and Methods

### Software

Analyses were performed using R (version 4.0.3), R Studio (version 2022.07.2). The application was developed using RShiny (version 1.7.3).

### TraDIS library ciprofloxacin challenging and sequencing

An *E. coli* K12 TraDIS library was generated as previously described (38) and challenged with subinhibitory ciprofloxacin (40 µg/mL) in 10 mL of Mueller Hinton cation adjusted media. Genomic DNA was extracted using the DNeasy UltraClean Microbial Kit (Qiagen) according to the manufacturer’s instructions and was sequenced on an Illumina HiSeq2500 platform at the Wellcome Sanger Institute.

### Identifying chromosomal location bias

The data was run through the Bio::TraDIS pipeline (4) using SMALT mapping and a minimum read count of 10. To identify whether CLB was present, the log2FCs were plotted in genome order, with locus on the x-axis and log2FC on the y-axis.

### Generating normalized read counts and offsets

For each condition, the read counts were obtained, and the first 1000 genes were appended to the end of the file and the last 1000 to the beginning to mimic a circular genome for the median sliding window function. Read counts of zero were discounted from the dataset to remove their influence on the median. The medianFilter function from package FBN (version 1.5.1) was used to calculate a median for each locus based on an adjustable window size. We have found that a default window size of 500 is sufficient for many smoother trends, whilst sharps trends need to be computed with a smaller window size. A ratio for each point was calculated by dividing the locus’ median by the mean median of all loci. The normalized read count was obtained by dividing the raw read count by the ratio computed for each locus. The normalized counts were not used as a replacement for the raw read counts, instead, an offset dataset was created as outlined in **Eq 1**. An arbitrary value of 0.1 was added before logging the counts to prevent the undefined log of 0 occurring. Loci with raw read counts of less than 10 were removed from the analysis.

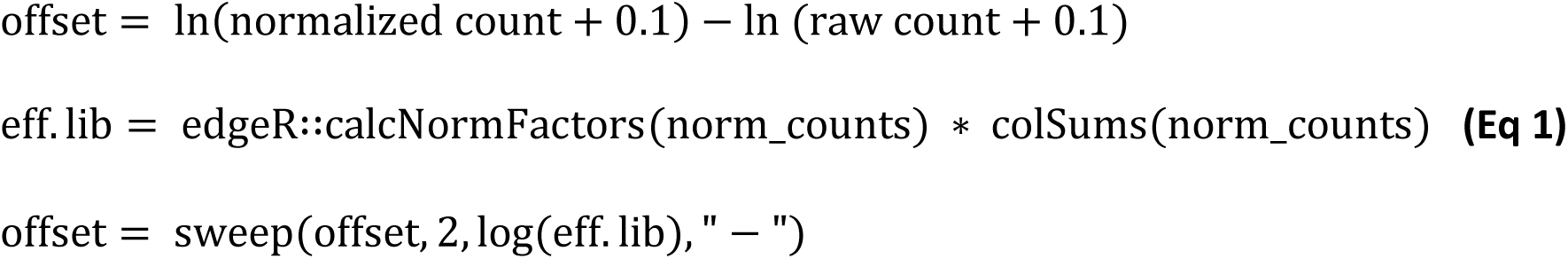

### Comparisons using edgeR

EdgeR (version 3.32.1) was used to compare the ciprofloxacin condition to the control to generate fold change data. Raw counts were put into a DGEList with groupings and scaleOffset() was used to offset the raw read counts. Library size normalization was not computed due to the inclusion of the offset, which produces a custom normalization factor per gene. Common negative binomial (estimateGLMCommonDisp()) and Bayes tagwise (estimateGLMTagwiseDisp()) dispersions for general linear models were calculated. A gene wise negative binomial for general linear models (glmFit() and glmLRT()) was fit with contrasts to produce likelihood ratio tests per gene between the control and conditions, producing the log fold change and adjusted p-values.

### Minimum inhibitory concentration assays

Minimum inhibitory concentration (MIC) assays were performed for the single gene knockouts to determine their breakpoint compared to wild type cultures. Mutants were steaked from frozen onto Mueller Hinton (MH) agar plates and incubated overnight at 37°C. Three single colonies of each mutant were inoculated into 5 mL of cation adjusted MH broth (CAMHB) and grown overnight at 37°C and 200 rpm shaking. A 1/100 dilution of the overnight cultures was made in 5 mL of fresh CAMHB and grown for 2.5 hours until exponential phase. MICs were performed with triplicate technical replicates in a 96-well plate with approximately 1×10^5^ cells per 150 µL well and grown overnight at 37°C with 200 rpm shaking. Cells were imaged after 16 hours at OD_600_ on a PHERAstar plate reader (BMG Labtech). Wells were blanked and averaged within triplicates. MIC was determined as the lowest concentration that inhibited at least 50 % of growth compared to the untreated mutant positive control.

### Code availability

The ChromoCorrect code is publicly available and can be found on Github at https://github.com/BarquistLab/ChromoCorrect and as an online interface at https://thecainlab.shinyapps.io/ChromoCorrect/.

### Data availability

TraDIS sequencing reads were deposited in the European Nucleotide Archive (ENA) under study accession number PRJEB35059. RNA sequencing data in *S. pneumoniae* reported in Slager *et al.* (25) sourced from ENA study accession PRJNA235855. RNA-seq data in *P. aeruginosa* reported in Murray *et al*. (21) sourced from ENA study accession PRJNA291292.

## Acknowledgements

The authors would like to thank Julian Parkhill and the Wellcome Sanger Institute sequencing team for kindly providing the TIS sequencing data used in this publication. We would also like to thank Claire Maher, Hannah Lott and Natasha Delgado for prototype testing of ChromoCorrect.

GJS acknowledges the Helmholtz Information & Data Science Academy (HIDA) for providing financial support for a short-term research visit to the Helmholtz Institute for RNA-based Infection Research (HIRI) and acknowledges an Australian Research Council-funded scholarship from project grant DE180100929. This project was partially funded by the Bavarian State Ministry for Science and the Arts through the research network <u>bayresq.net</u> to LB and Australian National Health and Medical Research Council (NHMRC) Project Grant 1159752 to AKC and LB. AKC was supported by an Australian Research Council (ARC) Future fellowship (FT220100152).

